# A Mechanism for Beam-Induced Motion in Cryo-Electron Microscopy

**DOI:** 10.1101/845552

**Authors:** Robert E. Thorne

## Abstract

Estimates of heat transfer rates during plunge cooling and patterns of ice observed in cryo-EM samples indicate that the grid bars cool much more slowly than do the support foil and sample near the middle of the grid openings. The resulting transient temperature differences generate transient tensile stresses in the support foil. Most of this foil stress develops while the sample is liquid and cooling toward its glass transition *T*_g_, and so does not generate tensile sample stress. As the grid bars continue cooling toward the cryogen temperature and contracting, the tensile stress in the foil is released, placing the sample in compressive stress. Radiation-induced creep in the presence of this compressive stress should generate a doming of the sample in the foil openings, as is observed experimentally. Crude estimates of the magnitude of doming that may be generated by this mechanism are consistent with observation. Several approaches to reducing beam-induced motion are discussed.

## 1. Introduction

In biomolecular single-particle cryo-electron microscopy, biomolecules in aqueous solution are deposited and then blotted or wicked to form a thin film on a thin holey carbon or metal support foil, that is in turn supported by a much thicker metal grid (Cheng *et al.*, 2015). These samples are rapidly cooled by plunging in liquid ethane, so as to vitrify the biomolecule containing film. Upon irradiation by electrons, the biomolecules (and the amorphous ice in which they reside) within the foil’s holes are observed to move (Glaeser, 2016; Vinothkumar & Henderson, 2016). This motion causes blurring of acquired images (Glaeser *et al.*, 2011; Brilot *et al.*, 2012; Russo & Passmore, 2016*b*,*a*; Glaeser, 2016; Henderson, 2018; Shi *et al.*, 2019). High efficiency, high frame rate direct electron detectors have allowed movies of this motion to be acquired (Campbell *et al.*, 2012; Faruqi & McMullan, 2018), and algorithms have been developed to model and correct for the effects of this motion (Li *et al.*, 2013; Rubinstein & Brubaker, 2015; Ripstein & Rubinstein, 2016; Zheng *et al.*, 2017). These advances have contributed to dramatic improvements in achievable resolution in cryo-EM, and to an explosion of interest in this approach to biomolecular structure determination.

However, beam-induced sample motion remains a serious factor limiting achievable resolution and the interpretation of acquired images. Experimental studies to date suggest the following salient features. First, motion is a function of dose, not time. Second, the rate of change of particle positions with dose is largest at low doses, below 2-4 e^−^/Å^2^ (Vinothkumar & Henderson, 2016), when accumulated radiation damage is modest and the highest-resolution structural information is available. The direction and magnitude of motion varies with position in each hole and between holes, but the largest motion component corresponds to a doming of the vitrified sample within the hole (Wright *et al.*, 2006; Brilot *et al.*, 2012; Campbell *et al.*, 2012; Zheng *et al.*, 2017). The amount of motion appears to be reduced by reducing the size of the holes in the foil (Russo & Passmore, 2016*b*).

Electron irradiation breaks bonds and increases the average distance between atoms, so that the sample film is expected to expand with dose (Vinothkumar & Henderson, 2016). The expansion should be roughly linear with dose (except at large doses where damage becomes severe), as is found in X-ray crystallography. This mechanism cannot explain the rapid initial motion but may account in part for more gradual motion at larger doses.

Electron irradiation may promote release via plastic creep of stress that is developed within the sample during cooling. Irradiation-induced stress relaxation has been extensively studied, e.g., in the context of materials properties for nuclear reactors (Bullough & Wood, 1980; Shibata, 2013). Creep is largest at the start of irradiation, when the stress is largest, and decreases as irradiation continues and the driving stress is relaxed by creep. This mechanism provides a natural explanation for the initial rapid beam-induced motion in single-particle cryo-EM samples. However, the nature and origin of the driving stress has not been identified.

Here we discuss a possible origin for cooling-induced stress in cryo-EM samples that is qualitatively consistent with observation, and how this stress may be reduced.

## 2. Mechanisms for sample stress generation

### 2.1 Cooling a biomolecule-containing sample on a freely sliding support foil generates tensile sample stress

Suppose that a biomolecule-containing sample is deposited on a holey foil, and that the foil is free to slip relative to the supporting grid. In this case, contraction of the foil during cooling is not coupled with that of the grid. Consequently, as the foil cools, it will contract at a temperature-dependent rate that depends only on its composition and its interaction with the sample.

The behaviour of the biomolecule-containing sample on cooling is more complex. Between room temperature and water’s glass transition temperature *T*_g,water_ ≈136 K (Amann-Winkel *et al.*, 2016), water undergoes a net volume expansion of ~8%. Since the sample is liquid in this temperature range, this expansion is essentially uncoupled from the contraction of the support foil. Consequently, differences in thermal contraction of the sample and foil between room temperature and *T*_g_ should generate no sample stress^1^.

On cooling from *T*_g_ to the final temperature *T*_cryo_ (typically ~90 K), the thermal contraction of bulk amorphous ice roughly matches that of hexagonal ice I_h_, with an average thermal expansion coefficient over this temperature range of 1.4 × 10^−5^ (Röttger *et al.*, 1994). Average thermal expansion coefficients over the same temperature range are smaller for all metals and other materials (amorphous carbon, Si, SiN) used as sample supports in cryo-EM; the average expansion coefficients of Au (~1.2 × 10^−5^) and Cu (~1.1 × 10^−5^) (Corruccini & Gniewek, 1961) provide the closest match to that of I_h_ in this temperature range. Thus, differences in thermal expansion coefficients between the sample and a freely sliding support foil should result in tensile sample stress once both have cooled to *T*_cryo_. Beam-induced sample creep would then produce thinning, rather than the observed doming, of the sample within the foil’s holes.

### 2.2 The sample support foil and grid are tightly coupled

In fact, the foil is in general not free to slip on the grid, but is tightly fixed to it by dispersion and electrostatic forces; by surface tension forces from sample that flows through the foil’s holes and wets the grid bars; and (once sufficiently cold) by vitrified or crystalline ice.

### 2.3 When the foil is in tension, its dimensions must match those of the grid

If the grid is stretched, the foil must stretch with it (at least until the foil’s tension reaches the limit of static friction between foil and grid). If the grid is compressed, the foil will buckle where it is not in contact with the grid. The grid is much thicker than the foil (~10 μm vs ~0.05 μm), and so has much greater elastic stiffness. Consequently, the thermal contraction of the grid material alone will (to first approximation) determine the foil’s dimensions at all temperatures, as long as the foil is in tension. Tension and buckling of the foil at the final temperature *T*_cryo_ can be minimized by using the same material (e.g., Au) for the grid and foil (Russo & Passmore, 2016a).

### 2.4 If grid, foil, and sample had the same temperature during cooling, differences in thermal expansion coefficients of grid, foil, and sample would determine sample stress

If the grid material has a larger average thermal expansion coefficient between *T*_cryo_ and 300 K than the foil, on cooling to *T*_cryo_ the foil (and sample) will buckle. If the grid has a smaller average thermal expansion coefficient, the foil’s physical contraction will match that of the grid, and the foil will develop tension. The stress developed in the sample will then be determined by the mismatch between its average thermal expansion coefficient between *T*_g_ and *T*_cryo_ and that of the grid (not the foil). Pure amorphous ice has a larger average expansion coefficient in this temperature range than all materials used in cryo-EM grids. Consequently, if grid, foil, and sample all had the same temperature during cooling, the sample would always be under tensile stress, and no doming upon irradiation would be expected.

### 2.5 During plunge cooling, grids cool more slowly that the support foil and biomolecule-containing sample

Pure water vitrifies when cooled at rates in excess of ~250,000 K/s (Warkentin et al., 2013). Routine observation of crystalline ice on and near the grid bars even when the sample on the foil away from the grid bars is fully vitrified provides direct evidence that grid bars cool more slowly.

In the boundary layer approximation for forced convective heat transfer, the coefficient of heat transfer *h* from the grid+foil+sample to the liquid cryogen depends only on the plunge speed, the grid dimensions, and the properties of the liquid cryogen (Kriminski *et al.*, 2002). The rate of heat transfer per unit area of foil and grid depends only on *h* and on the difference between the local grid/foil/sample temperature *T* and *T*_cryo_, *dq*/*dt* = *h* (*T* − *T*_*cryo*_). If the grid and foil were thermally isolated from each other, the cooling rates of each would then be proportional to their heat capacity per unit area. For a 50 nm gold foil on a 10 μm thick gold grid, these differ by a factor of ~10^3^.

Thermal conduction through the foil + sample from the grid bars to the center of the foil in each grid opening (as well as heat transfer from the grid to the liquid cryogen that then flows over the adjacent foil) will reduce the foil + sample’s cooling rate toward that of the grid. With non-metal (e.g., amorphous carbon) foils, the foil + sample’s thermal conductance *κ* is small, and the rate of heat transfer from grid bars through the foil per unit area (*dq*/*dt* ~ (*κt* / *w*^2^)(*T*_*grid*_ − *T*_*foil*_), where *w* is the width of a grid square and *t* is the foil thickness, will always be small compared with the rate of heat transfer from sample+foil to the liquid cryogen. With (high thermal conductivity) gold foils, the rates of heat transfer per unit temperature difference from grid to foil and from foil to liquid cryogen are more nearly comparable, i.e., *h* ~ *κt* / *w*^2^. However, this still implies that large temperature differences between the grid bars and the sample+foil near the center of each grid opening (and large temperature gradients within the sample+foil) must transiently occur during cooling, and that the grid bars will reach the final temperature *T*_cryo_ long after the sample+foil near the center of each grid opening.

Thermal mass near grid bars is further increased by the common presence of excess unwicked sample that flows through foil holes and wets to the back side of the foil and to the grid bars. Between room temperature and 90 K, the total heat capacity per unit *volume* of water/hexagonal ice is roughly 50% more than that of Au, and so excess sample on the grid bars can substantially slow their cooling. Ice is a poor thermal conductor, having a conductivity ~10^−2^ that of Au, and so has only a modest effect on thermal conduction from grid bars to the foil center.

### 2.6 Slower cooling of the grid produces transient tensile stress in the support foil

As shown in Figures 1 and 2 slower cooling of the grid bars than the foil+sample between them must produce transient tensile stress in the foil. The foil tries to contract toward the equilibrium length appropriate to its temperature, but is prevented from doing so by the more rigid grid that has cooled and contracted less. This is true even if the grid and foil are of the same material. The biomolecule-containing sample then vitrifies on a foil that is already under tensile stress. If the cooling rate of the grid bars is much smaller than that of the sample+foil between them, the temperature difference between grid bars and foil may continue to grow, the tensile stress in the foil may grow, and the sample may also develop a small amount of tensile stress as it cools from *T*_g_ to *T*_cryo_.

**Figure 1.**
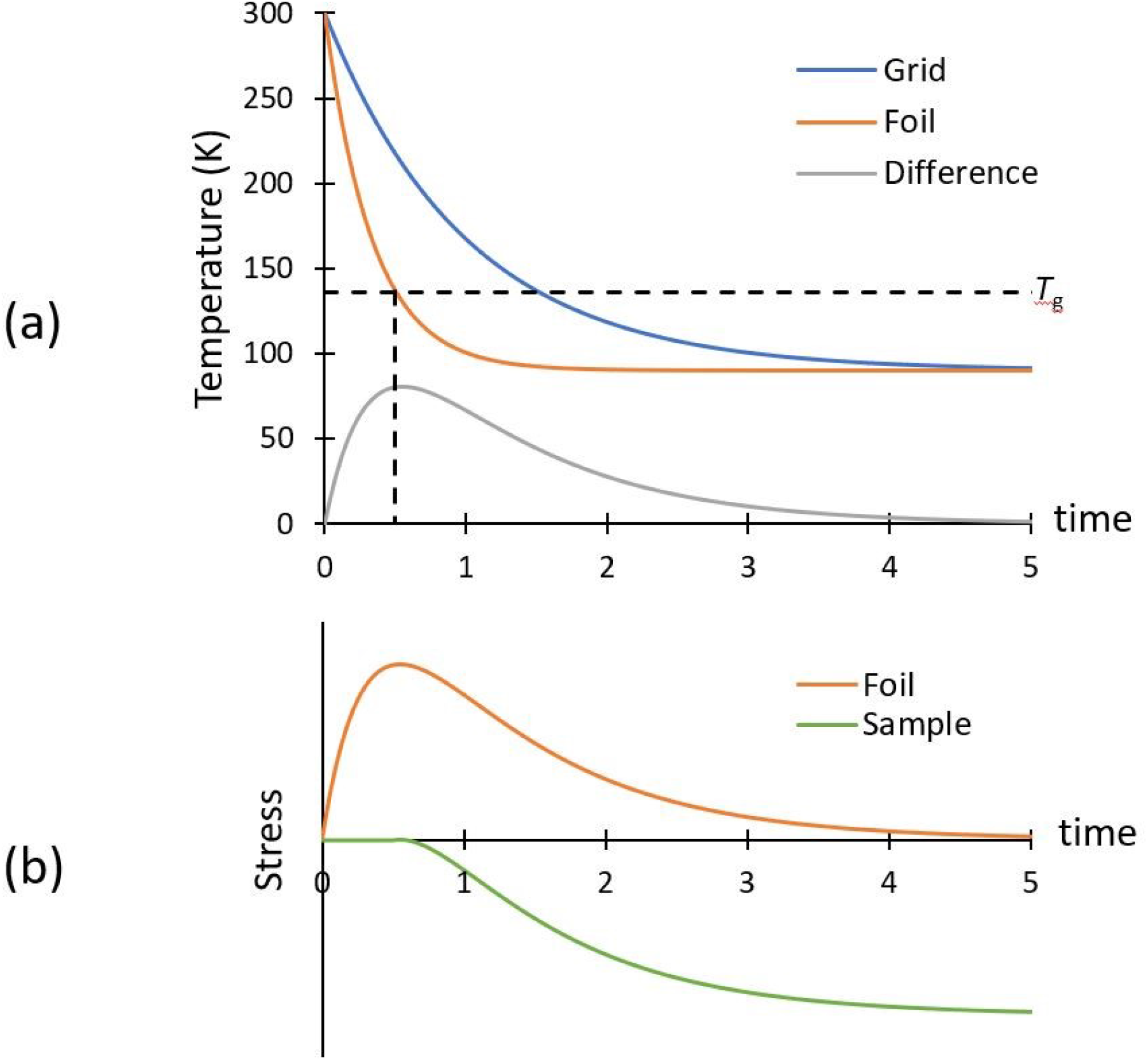
Temperature and stress during plunge cooling of cryo-EM samples. (a) The grid cools more slowly than the foil ands sample near the middle of the grid openings, so a large temperature difference between grid and foil may transiently occur. (b) The transient temperature difference produces a transient tensile stress in the foil, whose dimensions are constrained by those of the grid. The sample vitrifies and becomes strongly coupled to the foil only at *T*_g_, when the foil is under tensile stress. As the grid bars cool, the temperature difference between grid bars and foil decreases, and the tensile stress in the foil is released, the sample is placed under compressive stress. The grid and foil are here assumed to be of the same material.

**Figure 2.**
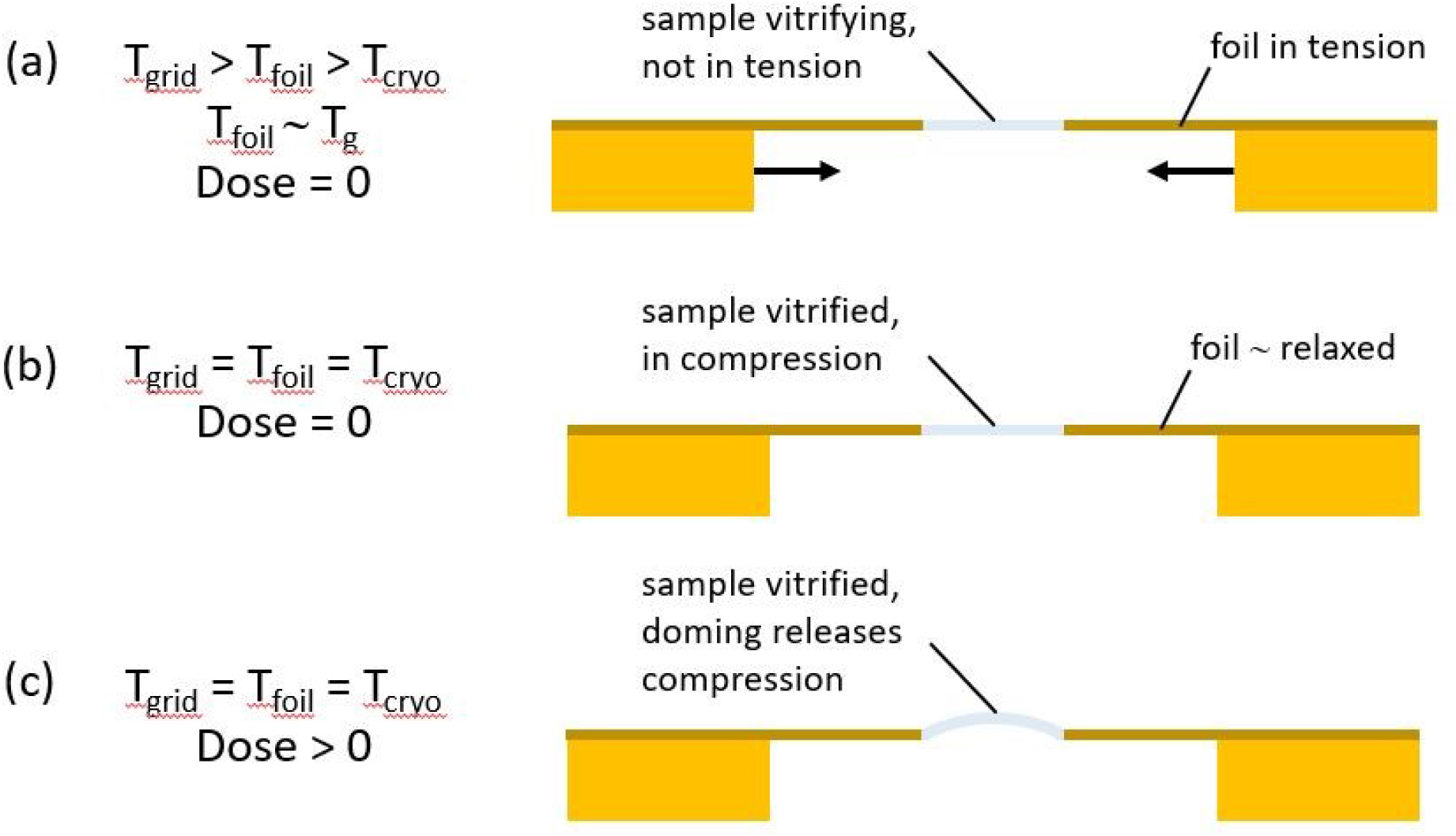
Schematic illustration of how compressive stress and doming may be generated during plunge cooling of cryo-EM samples. (a) Sample and foil have cooled to *T*_g_, and the foil is under tensile stress between the warmer grid bars. (b) Sample, foil, and grid have reached the final temperature *T*_cryo_; tensile stress in the foil associated with the transient temperature difference between grid and foil has been released, and the sample is now under compression. (c) After receiving an electron dose *D*, radiation induced creep in the presence of compressive stress has produced doming of the sample within the foil’s holes.

### 2.7 Transient tensile stress in the foil is released as the temperature of grid and foil converge to the final *T*_cryo_

As the grid continues to cool toward the final temperature *T*_cryo_ of grid and foil, the temperature difference and the difference in equilibrium length of grid and foil eventually decreases, and tensile stress in the foil decreases. If grid and foil are of the same material, the foil’s stress (in the absence of sample) will be completely released once they both reach *T*_cryo_.

### 2.8 Release of transient tensile foil stress places the sample under compressive stress

The sample vitrifies on a foil that is under tensile stress because the foil is colder than the grid bars. As the temperature difference between grid bars and foil goes to zero, the contracting grid allows the foil to contract toward its equilibrium length, and the foil stress is released. As the foil attempts to contract, this places the vitrified sample upon it under compressive stress.

### 2.9 Radiation-induced sample creep in the presence of this compressive stress may generate doming of the sample within the holes in the foil

When stressed (loaded), irradiated materials undergo creep in a way that releases the stress (Bullough & Wood, 1980; Shibata, 2013). In a simple model, the irradiation induced creep strain *ε*_*c*_ is given by *ε*_*c*_ = *aσ*(1− exp(−*bD*))+*cσD* where *D* is the radiation dose, σ is the stress, and *a*, *b*, and *c* are constants. In the presence of compressive sample stress parallel to the plane of the foil, the sample may release that stress by increasing its area via doming motion, as occurs when, e.g., a ruler is squeezed at its ends along its length.

For samples within foil holes of diameter 1.2 μm, Brillot et al. (2012) observed radiation-induced doming displacement of the sample, with a displacement of the central region perpendicular to the plane of the foil of ~150 Å after a dose of 32 e^−^Å^2^. The ratio of the surface area of a spherical cap of radius *a* and height *h* to the area of a disk of the same radius is 1+*h*^2^ / *a*^2^, and so the observed doming corresponds to a ~0.06% increase in sample area. Crudely, we can assume that this area increase is comparable to the fractional decrease in hole area from when the sample vitrifies to when the grid reaches the same final temperature *T*_cryo_ as the sample+foil. If the grid and foil are both made of Au, which has an average areal expansion coefficient of ~2.5×10^−5^ between room temperature and 90 K (Corruccini & Gniewek, 1961), then this change in hole area would be associated with a modest 25 K temperature difference between grid and sample + foil (near the grid opening centers) at the time of sample vitrification. The actual temperature differences could be as large as ~160 K, and this analysis neglects any contribution to doming within the holes from radiation induced creep of compressively stressed sample outside the holes. Consequently, both the sign and magnitude of the observed sample motion appear to be consistent with the present model.

## 3. Reducing beam induced motion

Based on the mechanism for rapid initial beam-induced motion discussed here, how might this motion be reduced?

First, the support foil and grid can be designed so that the foil can slip relative to the grid (at least in some regions of the grid), releasing tensile stresses that arise due to slower cooling of the grid than the foil. The limit of static friction between foil and grid will depend on the choice of foil and grid materials, their surface roughnesses, and their surface cleanliness/oxidation state. Inert layered materials like graphite might yield suitably low friction. Using a continuous rather than holey foil or a foil covered with thin graphene film may also help by preventing wetting of the sample to the grid bars and gluing of the grid and foil by vitrified or crystallized sample. This may in part explain recent reports of reduced sample motion using graphene-covered sample supports (Naydenova *et al.*, 2019).

Second, as already demonstrated (Russo & Passmore, 2016*a*), the hole size in the supporting foil can be reduced. Beam-induced sample creep in the presence of stress is likely constrained by contact with the supporting foil, so creep (and certainly doming) may be largely confined to the holes. In the simplest model, release of a given amount of stress Δ*σ* (an intensive quantity) will produce a fractional change in linear dimension Δ*L* / *L* and in area Δ*A* / *A* ∝ (Δ*L* / *L*)^2^. When a flat disk of radius *a* expands to a spherical cap of height *h* and the same base radius, its surface area changes by Δ*A* / *A* = (*h* / *a*)^2^. Consequently, *h* / *a* ∝ Δ*σ*, and the dome height will decrease with decreasing hole size.

Third, the grid can be made of a material that undergoes only modest thermal contraction on cooling. Mo, W, and doped Si have thermal expansion coefficients roughly 1/3, 1/3, and 1/15, respectively, of that of Cu and Au. This will reduce the maximum foil stress due to transient temperature differences between foil and grid, and thus the compressive sample stress generated when their temperatures converge.

Fourth, temperature differences between the foil and grid during cooling can be reduced by reducing the difference in their cooling rates. This can be accomplished by reducing the heat capacity of the grid by using a grid material with a lower specific heat per unit volume, or by reducing the width and/or thickness of the grid bars. Regions of reduced grid thickness/width, where imaging will be performed, can be mechanically and thermally decoupled via “weak links” from surrounding regions needed to provide mechanical stiffness for handling.

Finally, the temperature to which the sample is initially cooled can be increased by raising the temperature of the liquid ethane. As the foil cools more rapidly and reaches the final temperature much sooner than the grid, this will decrease the maximum temperature difference between foil and grid, and reduce the release of tensile stress in the foil as the grid cools to the final temperature.

Pure water’s glass transition is at 136 K, and it undergoes devitrification on warming above ~150 K (Mayer & Hallbrucker, 1987; Hallbrucker & Mayer, 1987). To ensure that the cooling time to below *T*_g_ is sufficiently short to achieve vitrification, the final temperature must be well below *T*_g_; how much below will depend on the sample and foil thicknesses, and also on the maximum tolerable ice fraction in the sample to achieve high resolution imaging.

For aqueous solutions, the glass transition temperature increases and the critical cooling rate required to achieve vitrification decreases with increasing solute concentration (Warkentin *et al.*, 2013). Consequently, increasing the solute concentration should allow higher ethane temperatures to be used. The solutes can include salts present in protein buffers as well as cryoprotectants like glycerol. However, these reduce electron density contrast between the biomolecule and buffer (Tyree *et al.*, 2018), and so reduce measurement signal to noise. Solutes also include the biomolecules themselves. These tend to be less effective in suppressing ice formation on a per unit mass basis than salts or glycerol. However, if they can be highly concentrated without aggregation, including via evaporation after deposition on the grid, they could be quite effective.

Very recently, Shi et al.(Shi *et al.*, 2019) have reported improved cryo-EM image resolution for apoferritin at low doses, suggesting reduced beam-induced motion, using samples cooled in ethane at 163 K. These results are consistent with the mechanism for beam-induced motion discussed here. The very high temperatures used - well above *T*_g_ for water and dilute aqueous solutions - are unlikely to yield ice-free samples under most conditions. Shi et al. only report FFTs of real space images, which provide much less sensitive detection of crystalline ice than direct diffraction measurements, so the presence of ice in their samples cannot be ruled out. However, the particle densities in their images are very high, and may have been high enough to give nearly fully vitrified samples at such high temperatures.

1 This ignores the short timescale (~1 ms) of cooling and any viscoelastic response of the sample on this timescale above T_g_.

## References

Amann-Winkel, K., Böhmer, R., Fujara, F., Gainaru, C., Geil, B. & Loerting, T. (2016). Rev. Mod. Phys. 88, 1–20.

Brilot, A. F., Chen, J. Z., Cheng, A., Pan, J., Harrison, S. C., Potter, C. S., Carragher, B., Henderson, R. & Grigorieff, N. (2012). J. Struct. Biol. 177, 630–637.

Bullough, R. & Wood, M. H. (1980). J. Nucl. Mater. 90, 1–21.

Campbell, M. G., Cheng, A., Brilot, A. F., Moeller, A., Lyumkis, D., Veesler, D., Pan, J., Harrison, S. C., Potter, C. S., Carragher, B. & Grigorieff, N. (2012). Structure. 20, 1823–1828.

Cheng, Y., Grigorieff, N., Penczek, P. A. & Walz, T. (2015). Cell. 161, 438–449.

Corruccini, R. & Gniewek, J. (1961). Thermal expansion of technical solids at low temperatures.

Faruqi, A. R. & McMullan, G. (2018). Nucl. Inst. Meth. Phys. Res. A. 878, 180–190.

Glaeser, R. M. (2016). Methods in Enzymology, Vol. 579, pp. 19–50. Elsevier Inc.

Glaeser, R. M., McMullan, G., Faruqi, A. R. & Henderson, R. (2011). Ultramicroscopy. 111, 90–100.

Hallbrucker, A. & Mayer, E. (1987). J. Phys. Chem. 91, 503–505.

Henderson, R. (2018). Angew. Chemie - Int. Ed. 57, 10804–10825.

Kriminski, S., Caylor, C. L., Nonato, M. C., Finkelstein, K. D. & Thorne, R. E. (2002). Acta Cryst. D. 58, 459–471.

Li, X., Mooney, P., Zheng, S., Booth, C. R., Braunfeld, M. B., Gubbens, S., Agard, D. A. & Cheng, Y. (2013). Nat. Methods. 10, 584–590.

Mayer, E. & Hallbrucker, A. (1987). Nature. 325, 601–602.

Naydenova, K., Peet, M. J. & Russo, C. J. (2019). Proc. Natl. Acad. Sci. U. S. A. 116, 11718–11724.

Ripstein, Z. A. & Rubinstein, J. L. (2016). Methods in Enzymology, Vol. 579, pp. 103–124. Elsevier Inc.

Röttger, K., Endriss, A., Ihringer, J., Doyle, S. & Kuhs, W. F. (1994). Acta Cryst. B. 50, 644–648.

Rubinstein, J. L. & Brubaker, M. A. (2015). J. Struct. Biol. 192, 188–195.

Russo, C. J. & Passmore, L. A. (2016a). Curr. Opin. Struct. Biol. 37, 81–89.

Russo, C. J. & Passmore, L. A. (2016b). J. Struct. Biol. 193, 33–44.

Shi, H., Ling, W., Zhu, D. & Zhang, X. (2019). Increasing vitrification temperature improves the cryo-electron microscopy reconstruction. https://www.biorxiv.org/content/10.1101/824698v1

Shibata, T. (2013). Handbook of Advanced Ceramics, Vol. pp. 113–123.

Tyree, T. J., Dan, R. & Thorne, R. E. (2018). Acta Cryst. D. 74, 471–479.

Vinothkumar, K. R. & Henderson, R. (2016). Q. Rev. Biophys. 49,.

Warkentin, M. A., Sethna, J. P. & Thorne, R. E. (2013). Phys. Rev. Lett. 110, 015703.

Wright, E. R., Iancu, C. V., Tivol, W. F. & Jensen, G. J. (2006). J. Struct. Biol. 153, 241–252.

Zheng, S. Q., Palovcak, E., Armache, J. P., Verba, K. A., Cheng, Y. & Agard, D. A. (2017). Nat. Methods. 14, 331–332.

